# Aging Atlas Reveals Cell-Type-Specific Regulation of Pro-longevity Strategies

**DOI:** 10.1101/2023.02.28.530490

**Authors:** Shihong Max Gao, Yanyan Qi, Qinghao Zhang, Aaron S. Mohammed, Yi-Tang Lee, Youchen Guan, Hongjie Li, Yusi Fu, Meng C. Wang

**Affiliations:** Department of Molecular and Human Genetics, Baylor College of Medicine; Houston, TX 77030, USA; Program in Developmental Biology, Baylor College of Medicine; Houston, TX 77030, USA; Huffington Center on Aging, Baylor College of Medicine; Houston, TX 77030, USA; Janelia Research Campus, Howard Hughes Medical Institute; Ashburn, VA 20147, USA; Department of Biomedical Sciences, Creighton University School of Medicine; Omaha, NE 68178, USA

## Abstract

Organism aging occurs at the multicellular level; however, how pro-longevity mechanisms slow down aging in different cell types remains unclear. We generated single-cell transcriptomic atlases across the lifespan of *Caenorhabditis elegans* under different pro-longevity conditions (http://mengwanglab.org/atlas). We found cell-specific, age-related changes across somatic and germ cell types and developed transcriptomic aging clocks for different tissues. These clocks enabled us to determine tissue-specific aging-slowing effects of different pro-longevity mechanisms, and identify major cell types sensitive to these regulations. Additionally, we provided a systemic view of alternative polyadenylation events in different cell types, as well as their cell-type-specific changes during aging and under different pro-longevity conditions. Together, this study provides molecular insights into how aging occurs in different cell types and how they respond to pro-longevity strategies.

---

Aging in multicellular organisms involves functional declines in both somatic and reproductive tissues. However, how age-related molecular changes differ in various tissues at cellular resolution remains poorly understood. On the other hand, multiple pro-longevity strategies have been discovered in multicellular organisms ranging from *C. elegans* to mice, and some of them display high tissue-specificity in their regulations. For example, the reduction of insulin/IGF-1 signaling is a well-conserved mechanism in prolonging lifespan^1,2^ and in *C. elegans* and *Drosophila melanogaster*, fat storage tissues (intestine in worms and fat body in fruit flies) and neurons are two crucial sites for this pro-longevity mechanism^3,4^ Similarly, the reduction of TOR (target of rapamycin) signaling by either genetic or pharmacological interventions increases lifespan in a variety of organisms^5^; and in *C. elegans*, this longevity-promoting effect has been linked with neuronal or intestinal regulation^6–8^. In addition, our studies discovered that a lysosomal acid lipase LIPL-4 in the intestine of *C. elegans* induces specific lipid signals to activate nuclear transcription cell-autonomously and up-regulate the neuropeptide pathway cell non-autonomously, both leading to lifespan extension^9,10^. Despite their well-characterized roles in prolonging lifespan, whether and how these strategies slow aging of different tissues in distinct manners are yet to be determined.

## Generating the adult *C. elegans* cell atlas during aging

In recent years, single-cell and single-nucleus RNA sequencing (scRNA-seq and snRNA-seq) have proven to be effective ways to systemically profile transcriptomes at the single-cell resolution and have facilitated the discovery of cell-type-specific transcriptomic signatures in different tissues^11–19^. Recent studies showed that snRNA-seq is less biased for tissue sampling in atlas studies compared to scRNA-seq, because certain cell types (e.g., muscle and epidermal cells) cannot be efficiently isolated using single cell dissociation methods^20,21^. Thus, we have developed an snRNA-seq pipeline for systemically profiling transcriptomic changes in adult *C. elegans* at the single-cell resolution (Fig. 1A). For each condition, we harvested and homogenized ~2,000 worms. Nuclei were isolated using fluorescence-activated cell sorting (FACS) based on the DNA content signal (fig. S1A), and snRNA-seq was performed using the 10× Genomics platform. For each condition, 10,000 nuclei were sequenced to capture the transcriptome of 959 somatic cells and ~2,000 germ cells in adult *C. elegans*. After preprocessing and cell filtering, we generated 177,733 single-nuclei gene expression profiles. From this dataset, we were able to build an adult cell atlas that covers 15 major cell classes, including neurons, glia, hypodermis, intestine, muscle, pharynx, coelomocyte, gonadal sheath cells, vulva and uterus, uterine seam cells, distal tip cells and excretory gland cells, germline, sperms, spermatheca and embryonic cells (Fig. 1B and C). Note that adult *C. elegans* consists of various fully differentiated somatic cells and carries germ cells at different developmental stages and early embryos. Sub-clustering of these major cell classes further revealed many more different cell types for each class. For neurons, 71 subclusters were identified, which account for 104 of all 114 neuron classes in adult *C. elegans* (Fig. 1D)^22^. For the hypodermis, we further distinguished seam cells, rectal and vulval epithelium, and four tail tip hypodermal cells (hyp8-11) from other hypodermal cells (fig. S1B). For the muscle, we discovered four subclusters, including body wall muscle, head muscle, nonstriated muscle and vulva muscle (fig. S1C).

**Fig. 1.**
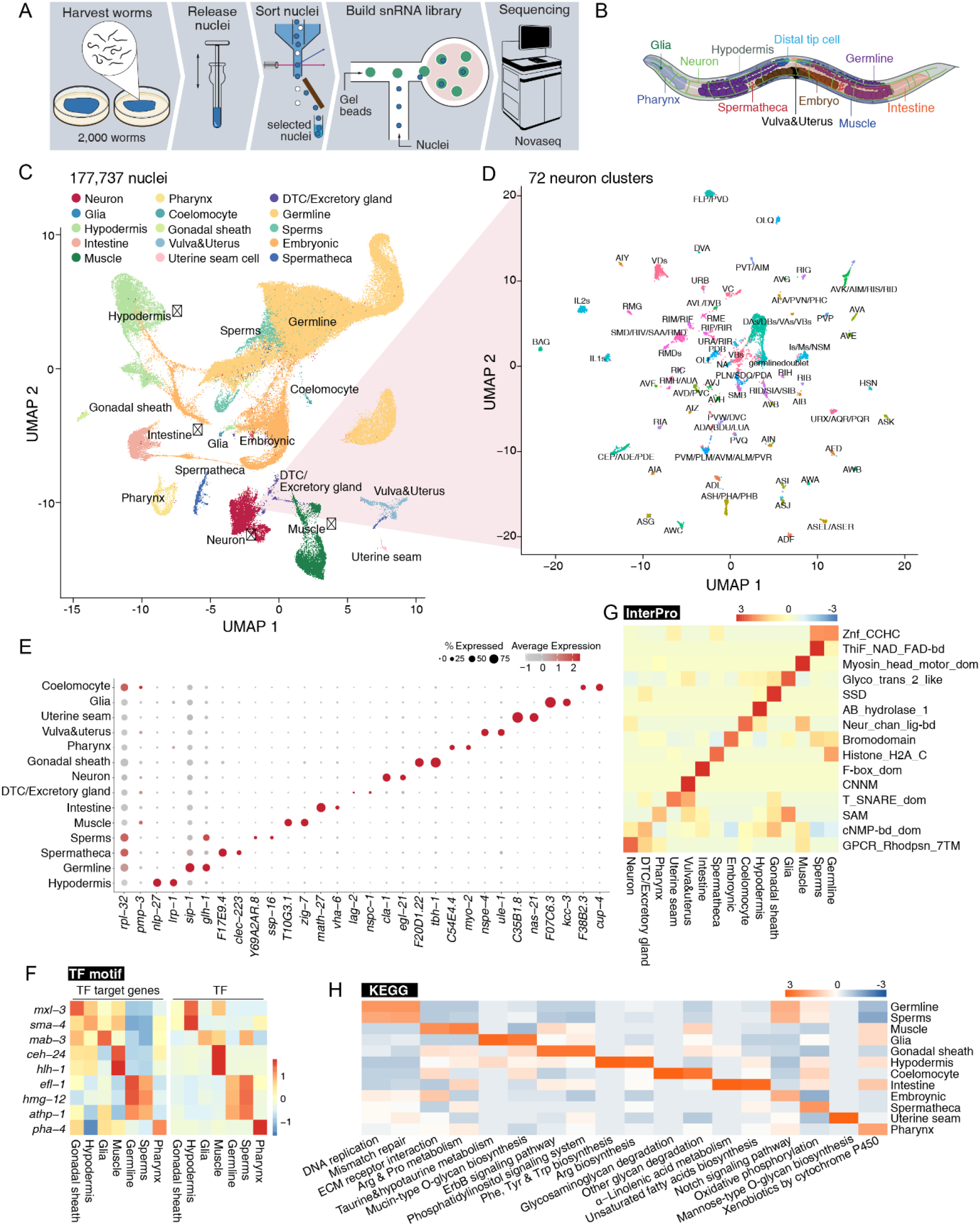
Adult cell atlas profiled at the single-cell resolution. (**A**) Schematics of single-cell transcriptome profiling in adult *C. elegans*. (**B**) Adult *C. elegans* anatomy with major tissues. (**C**) UMAP visualization of the single cells from adult *C. elegans* cell atlas, clusters corresponding to major tissues are shown, and tissues marked with * can be further subclustered. (**D**) Zoom-in 5 UMAP visualization of neuron sub-clustering into 72 subsets of neurons. (**E**) Dot plot showing the cell-type-specific expression pattern of known and newly identified markers for each major tissue. (**F**) Heatmaps showing the transcription factors (right) with the expression of their target genes (left) enriched explicitly in each tissue. (**G, H**) Heatmaps showing tissue-specific protein families based on InterPro (**G**) and KEGG pathways enrichment (**H**).

These large-scale profiling and clustering analyses revealed cell-type-specific transcriptional signatures that are supported by previously identified gene markers (Fig. 1E). Importantly, we identified several new cell-type-specific gene markers, which showed comparable or better specificity than well-known markers (Fig. 1E). We further conducted DNA binding motif analysis to search for transcription factors that could mediate cell-type-specific gene expression. We identified a number of transcription factors whose target genes are enriched in specific tissues, including *mxl-3* for gonad sheath cells, *sma-3* for the hypodermis, *mab-3* for glia, *ceh-24* and *hlh-1* for the muscle, *efl-1, hmg-12* and *atph-1* for germline/sperms, and *pha-4* for the pharynx (Fig. 1F). Accordingly, the expression of these transcription factors exhibited the same tissue specificity (Fig. 1F). Together, these snRNA-seq data sets provide a systemic overview of cell-type-specific transcriptional signatures and their regulations.

Next, we utilized InterPro and KEGG classification to analyze these cell-type-specific transcriptome profiles and discovered distinct functional features for each cell type. Some of these functional features are expected. For example, based on the annotation of InterPro classification, the G protein-coupled receptor family (GPCR_Rhodpsn_7TM) that represents crucial molecular sensors was enriched explicitly in neurons, and the myosin head motor domain family (Myosin_head_motor_dom) that is required for muscle contraction was specially enriched in the muscle (Fig. 1G). The gene module analysis for KEGG pathway enrichment revealed that alpha-linolenic acid metabolism and unsaturated fatty acid biosynthesis categories are enriched in the intestine, the major fat storage tissue of *C. elegans* (Fig. 1H). DNA replication and mismatch repair categories were enriched in the germline and sperms (Fig. 1H). We also uncovered several tissue-specific functional signatures that were not characterized before. For example, in glial cells, we discovered the specific enrichment of glycosyltransferase 2-like family (Glyco_trans_2_like) and mucin-type O-glycan biosynthesis based on annotation from InterPro (Fig. 1G) and KEGG (Fig. 1H), respectively, which together suggest the importance of O-glycosylation in glial physiology. InterPro and KEGG analyses also revealed that the α/β hydrolase superfamily (AB_hydrolase_1, Fig. 1G) and Arginine, Phenylamine, Tyrosine and Tryptophan biosynthesis (Fig. 1H) are specifically enriched in the hypodermis, indicating the active involvement of this tissue in metabolic processes.

Next, we focused on establishing the aging cell atlas to comprehensively understand age-related transcriptomic changes at the single-cell level. We first analyzed data from wild type (WT) worms at four different adult ages, day 1, day 6, day 12 and day 14 when the survival rate is 100%, 99%, 61% and 14%, respectively (Fig. 2A). We were able to construct aging cell atlases (fig. S2A-D). We found that the relative number of cells in different somatic tissue clusters remain the same during the aging process (fig. S2E), confirming that our snRNA-seq pipeline did not introduce sampling bias. However, we did observe the number of cells in the germline cluster decreases with aging (fig. S2F). Notably, we aged WT adult worms under normal physiological conditions, without interfering with their reproductive processes. This experimental setup enabled us to investigate transcriptomic changes during both somatic and reproductive aging.

**Fig. 2.**
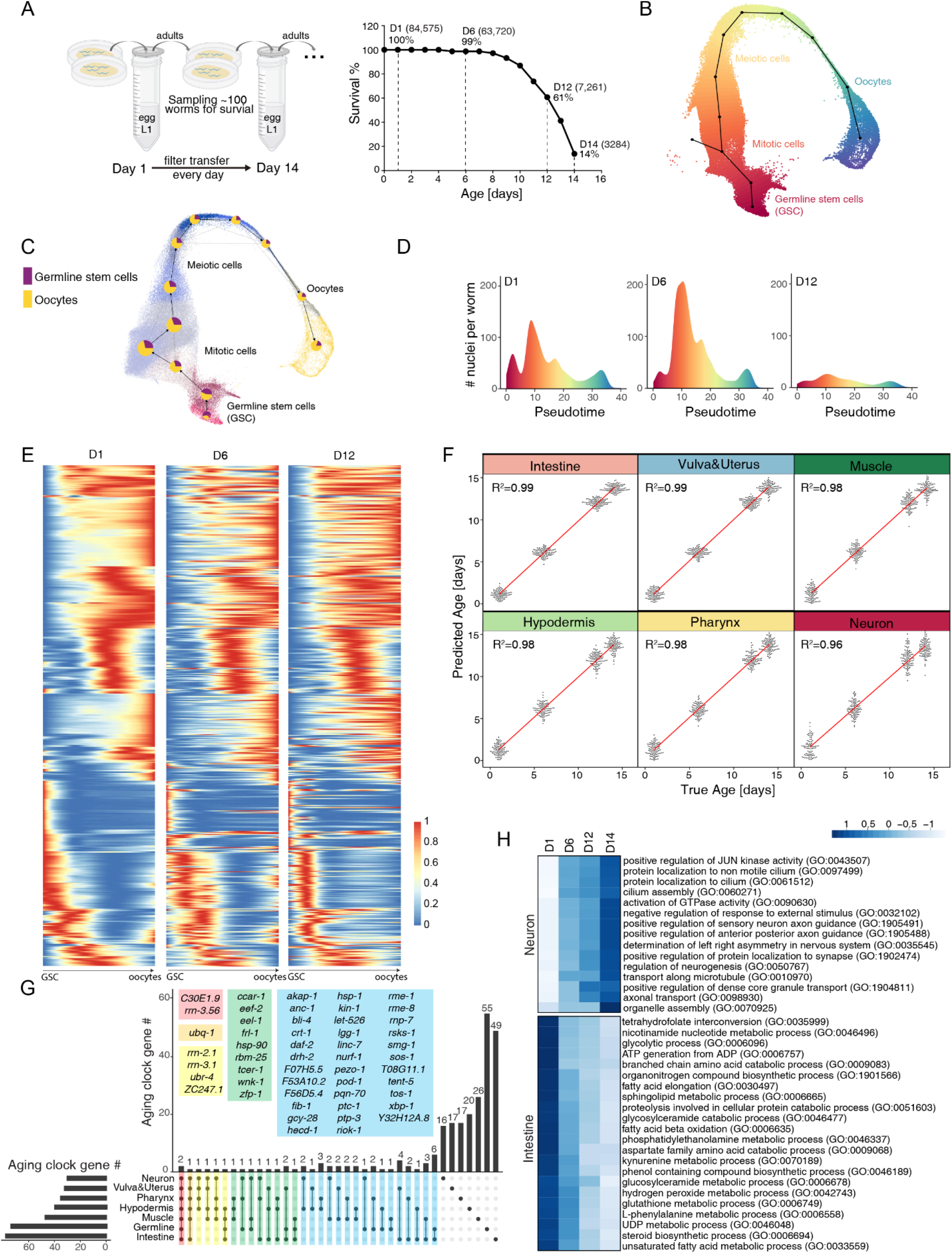
Aging cell atlas revealing cell type-specific transcriptomic changes during aging. (**A**) Schematic (left) of aging sample preparation under physiological conditions without interrupting worm reproduction. The survival curve (right) of worm samples used for nuclei collection at four time points to build the aging cell atlas. (**B**) Germ cell trajectory pseudotime presented in UMAP. (**C**) Germ cell trajectory PAGA map showing cell fate commitment. (**D**) Density plots showing the distribution of germline nuclei number along the pseudotime at different ages. (**E**) Heatmaps showing that gene expression temporal patterns along the developmental progression of germ cells were disrupted with aging. (**F**) Jitter plots showing the correlation between the true age and the predicted age from the age clock for each tissue. (**G**) UpSet plot showing the overlap between aging clock genes identified in each tissue. Genes identified in more than one tissuespecific aging clock are highlighted in colored boxes. (**H**) Heatmaps showing neuron- and intestine-specific GO terms enrichment changes during aging.

## Mapping germ cell fate trajectories during aging

To systematically understand transcriptomic changes in germ cells during aging, we generated germ cell fate trajectory maps and conducted pseudotime inference analyses. In order to minimize the impact of analysis bias, we utilized two different computational methods: the *Slingshot* package^23^ and RNA velocity-based scVelo algorithm^24^. The results obtained from these two methods were consistent with each other. These trajectory maps depicted the progression of germ cells, as they undergo proliferation and differentiation from germline stem cells, through mitotic cells and meiotic cells, towards mature oocytes (Fig 2B, C, fig. S3A). The unsupervised scVelo model computed the initial and terminal states within the trajectory and identified cell clusters at the root and the endpoints, which correspond to germline stem cells and mature oocytes, respectively (fig. S3B). Furthermore, directed partition-based graph abstraction (PAGA) provided a quantitative assessment of cell fate probabilities for the initial, intermediate and terminal states (Fig. 2C), consistent with the developmental order from germline stem cells to fully differentiated oocytes.

Importantly, the computational trajectory allowed us to construct a pseudotemporal order and analyze age-related changes in germ cells. We observed a drastic decrease in the number of germline stem cells as the worm aged from day 1 to day 12, while the number of germ cells in the mitotic-meiotic transition peaked at day 6 and decreased at day 12 (Fig. 2D). In contrast, the number of mature oocytes showed no changes between day 1 and day 6 but decreased at day 12 (Fig. 2D). These results suggest that various groups of germ cells undergo unique changes as organisms age, implying divergent aging processes within the germline.

To gain a molecular insight into these age-related changes, we identified hundreds of genes that display specific expression patterns within different germ cell groups and that show significant changes in their expression patterns during aging (top 500 genes highlighted in Fig. 2E and Table S1). These analyses reveal the molecular signatures for germ cells at various stages, and provide a comprehensive view of germline molecular alterations during aging.

## Tissue-specific aging clocks and functional analyses

Next, we leveraged tissue-specific transcriptomic changes during aging to build age-prediction models – aging clocks – for different tissues. Using machine learning, we constructed regression-based tissue-specific aging clocks that accurately predicted true chronological ages with correlations (R^2^) greater than 0.96 for tissues with more than 50 cells in the cluster (Fig. 2F, fig. S3C). Surprisingly, we found that those aging-clock genes are mostly tissue-specific and exhibit very little overlap between them (Fig. 2G), suggesting that different tissues may age differently with their unique transcriptional signatures.

To gain a more comprehensive understanding of functional changes associated with somatic aging in a tissue-specific manner, we have profiled cell-type-specific gene ontology (GO) term enrichment changes during aging. Our analysis discovered a series of GO terms whose enrichment levels in specific tissues showed a consistent trend of increasing or decreasing as age increases (age-related GO terms) (Fig. 2H, fig. S3D). Notably, we found that these age-related GO terms specific to neurons undergo expression increases from day 1 to day 14 (Fig. 2H). These include diverse gene categories crucial for neural functions, such as neurogenesis, dense core granule transport, axonal transport and guidance, and protein localization to the cilium and synapse (Fig. 2H). At the level of signal transduction, positive regulation of JUN kinase and GTPase activities also exhibited age-related increases in neurons (Fig. 2H). These changes suggest that neurons become hyperactivated during aging. Interestingly, neural hyperactivity has been linked with age-related neurodegeneration^25–27^ and lifespan reduction^28^; while suppressing neuronal hyperactivity extends lifespan^28^. In contrast, those age-related GO terms specific to the intestine showed age-related decreases in expression, including a variety of metabolic pathways, such as ATP production, branched chain amino acid catabolism, fatty acid unsaturation, elongation and oxidation, sphingolipid metabolism, and hydrogen peroxide metabolism (Fig. 2H). The intestine is a major metabolic tissue in *C. elegans*, and our snRNA-seq analysis provides a systemic, molecular understanding of its functional decline during aging.

## Aging clocks predicting pro-longevity effects in different tissues

Multiple strategies have been discovered that extend lifespan in various organisms ranging from *C. elegans* to mice. How do different pro-longevity mechanisms affect age-related transcriptomic changes in specific tissues? To address this question, we performed snRNA-seq analysis on three different long-lived strains at two different ages (day 1 and day 6 when survival rates are close to 100%, to focus on early changes that could contribute to longevity and minimize the impact of confounding factors related to organismal death). The *lipl-4* transgenic strain (*lipl-4 Tg*) constitutively expresses the LIPL-4 lysosomal acid lipase in the intestine, which extends lifespan by 40%-60% (Fig. 3A)^9,29^ DAF-2 is the *C. elegans* homolog of the insulin/IGF-1 receptor, and its loss-of-function (lf) mutant doubles the lifespan (Fig. 3B)^1,30^. The *rsks-1 gene* encodes S6 kinase in *C. elegans and* the *rsks-1(lf*) mutant reduces TOR signaling and shows 20-30% lifespan extension (Fig. 3C)^31^. We systematically profiled genes that showed differential expression between WT and long-lived strains in different cell clusters (Fig. 3D, 3E, Table S2), revealing tissue-specific transcriptome signatures that underscore the distinct pro-longevity mechanisms at the single-cell level. When compared to WT worms, we observed that the three long-lived strains exhibit the largest difference in the number of differentially expressed genes (DEGs) in the hypodermis (Fig. 3D, 3E), which can be attributed to the highest proportion of hypodermal cells among the recovered somatic nuclei (~30%, fig. S2E). The second largest DEG difference was found in neurons, where we observed substantial transcriptome differences in the *daf-2(lf*) and the *rsks-1(lf*) strains compared to WT at day 1. Notably, these differences were significantly reduced at day 6 (Fig. 3D, 3E). On the other hand, the *lipl-4 Tg* strain exhibited minor changes in the neural transcriptome on day 1 but displayed a large difference on day 6 (Fig. 3D, 3E). These results suggest that diverse tissues are affected by different pro-longevity mechanisms in distinct ways during the aging process.

**Fig. 3.**
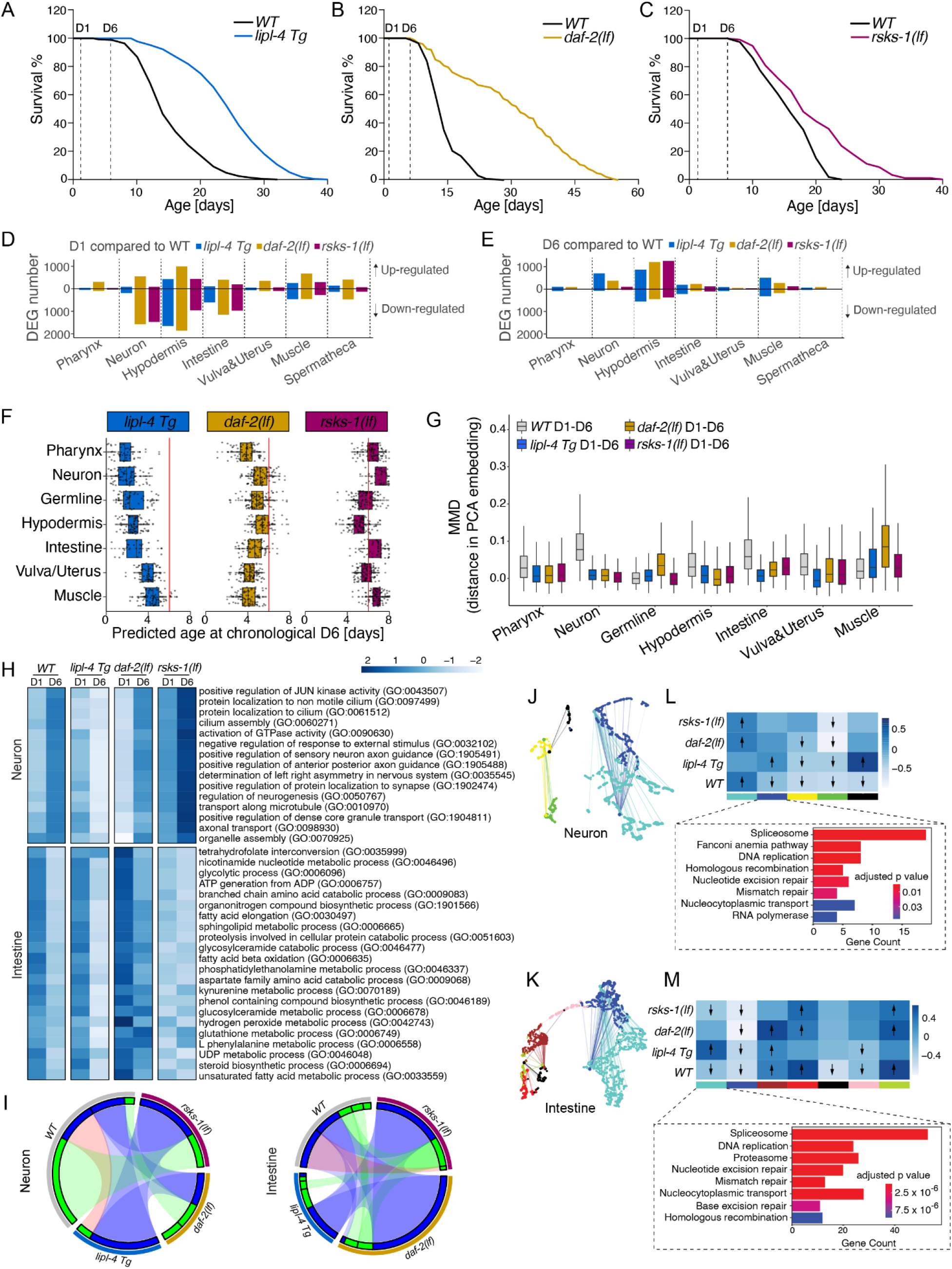
Tissue-specific regulation by different pro-longevity mechanisms. (**A-C**) Lifespans of long-lived *lipl-4* transgenic strain (*lipl-4 Tg*) (**A**), *daf-2* loss-of-function mutant (*daf-2(lf*)) (**B**), and *rsks-1* loss-of-function mutant (*rsks-1(lf*)) (**C**) compared to WT. (**D, E**) Bar graph showing the difference in the number of DEGs in major tissues between WT and long-lived strains at day 1 (**D**) and day 6 (**E**). Bars above or below zero representing higher or lower expression in long-lived worms than WT, respectively. (**F**) Boxplots showing the predicted biological ages of different tissues in three long-lived strains at the chronological age of day 6 (red lines) with tissue-specific aging clocks. (**G**) Boxplots displaying maximum mean discrepancy (MMD) between cells from day 1 and day 6 across different tissues, for both WT and long-lived worms. (**H**) Heatmaps showing the enrichment change of neuron- and intestine-specific GO terms between day 1 and day 6 in WT and long-lived strains. (**I**) Circos plots showing conserved co-expression modules (Fisher’s exact test, *P* < 0.01) that were significantly correlated with aging (Pearson’s correlation, *P* < 0.001 and R^2^ > 0.2) in neurons (left) and the intestine (right) between different genotypes. Blue ribbons connected conserved models that were both negatively correlated with aging in different genotypes, green ribbons connected conserved models that were both positively correlated with aging, and red ribbons connected conserved models that were oppositely correlated with aging in different genotypes. (**J, K**) UMAP visualization of the consensus co-expression network for aging-related modules (Pearson’s correlation, *P* < 0.001 and R^2^ > 0.2) in neurons (**J**) and the intestine (**K**). Dots represented genes and were colored by the module they belonged to. Edges represented co-expression between genes. (**L, M**) Correlations between consensus co-expression modules with aging, with significant modules (Pearson’s correlation, *P* < 0.001 and R^2^ > 0.2) marked by arrows representing the way they correlated with aging. Bar graphs in dashed boxes showed the KEGG pathway enrichment analysis (Fisher’s exact test, BH adjusted *P* < 0.01) for the blue module in neurons (**L**) and the turquoise module in the intestine (**M**).

To quantitatively assess the effects of these pro-longevity mechanisms on tissue-specific aging processes, we utilized the tissue-specific aging clocks trained in WT worms to predict the biological ages of different tissues in these long-lived strains at the chronological age of day 6. Our analysis revealed that both the *lipl-4 Tg* and *daf-2(lf*) strains exhibit younger predicted ages for all tissues than day 6, ranging from day 2 to day 5 (Fig. 3F). However, in the *rsks-1(lf*) strain, only hypodermal cells showed a predicted age younger than day 6 (Fig. 3F). Interestingly, while the *daf-2(lf*) strain has a stronger lifespan extension than the *lipl-4 Tg* strain, the latter showed a stronger effect on slowing down the aging clocks in different tissues (Fig. 3F). In parallel, we calculated the distance drift in the transcriptome of different tissues from day 1 to day 6 in WT and the long-lived worms using scMMD (maximum mean discrepancy) method. We found that in WT worms, neurons exhibited the biggest transcriptome drift from day 1 to day 6, followed by intestinal cells, which were largely attenuated in the three long-lived strains (Fig. 3G). These results suggest that these two tissues are more sensitive to aging than the other somatic tissues and play crucial roles in the endocrine regulation of longevity, as supported by previous genetic studies^32^ Furthermore, these results highlight the diverse impact of the three pro-longevity mechanisms on tissue-specific aging processes.

## Molecular regulation of tissue aging by different pro-longevity mechanisms

At the functional level, we found that the *rsks-1(lf) and* the *daf-2(lf*) mutant strains showed similar increases for the neuron-specific, age-related GO terms, like those observed in WT worms (Fig. 2H). In contrast, the *lipl-4 Tg* strain presented an opposite decreasing trend (Fig. 3H), which is in accordance with its strong aging-slowing effect in neurons based on the tissuespecific aging clocks (Fig. 3F). Meanwhile, we found that different pro-longevity mechanisms selectively suppress the decrease in intestine-specific, age-related GO terms. For example, tetrahydrofolate interconversion and UDP metabolic process did not show age-related decrease only in the *lipl-4 Tg* strain, while the glycosylceramide, kynurenine and L-phenylalanine processes remained unchanged during aging in the *daf-2(lf*) strain but were increased in the *rsks-1(lf*) strain. Additionally, the *rsks-1(lf*) strain specifically reversed the age-related decrease in the steroid biosynthesis process (Fig. 3H). These findings reveal the varying impact of distinct prolongevity mechanisms on tissue functions during the aging process.

To gain a systematic understanding of the molecular-level tissue-specific regulations, we further performed weighted gene co-expression network analysis (WGCNA)^33^. Initially, we constructed consensus co-expression networks for each tissue and identified co-expression modules that showed significant correlation with aging, either negatively (in blue) or positively (in green), in different genotypes including WT and three long-lived strains (Fig. 3I, fig S4A). We observed that modules negatively correlated with aging were preserved across genotypes in both neural and intestinal cells, while modules positively correlated with aging tended to be genotype-specific (Fig. 3I). Furthermore, we constructed consensus co-expression networks that reveal the co-expression connections between genes in each specific tissue (Fig. 3J, 3K, fig S4B) and demonstrated how the co-expression modules change with aging in different genotypes and tissues (Fig. 3L, 3M, fig S4C). Interestingly, we found that in neural cells, the blue module exhibits an age-related decrease in WT, but the decrease was absent in all three long-lived strains (Fig. 3L). KEGG pathway analysis further revealed the enrichment of the spliceosome in this module (Fig. 3L). Similarly, in intestinal cells, we observed the enrichment of the spliceosome in the turquoise module, which was decreased in both WT and the *rsks-1(lf*) strain, but not in the *daf-2(lf*) strain or the *lipl-4 Tg* strain (Fig. 3M). Together, these findings indicate that the splicing pathway may be involved in the regulation of aging and longevity, in a cell-type and genotype specific manner.

## Age-related APA changes and their regulation by pro-longevity pathways

Alternative splicing and alternative polyadenylation (APA) are two critical RNA-processing mechanisms that are frequently interconnected. While alternative splicing has been linked to longevity regulation in a variety of organisms, including *C. elegans*^34^, it is currently unclear whether APA undergoes age-related changes in different cell types and whether pro-longevity mechanisms influence APA. To obtain a systemic view of APA changes during aging in different cell types, we utilized the polyApipe tool to calculate the type of APA of pre-mRNAs (Fig. 4A) at the single-cell resolution in various cell clusters. Our analysis identified 851 candidate genes that showed a tissue-specific preference in their use of APA sites (Method, Table S3). Subsequently, we selected 55 genes that were highly expressed in over 20% of cells (Fig. 4B). Interestingly, germline cells and sperms exhibited a clear preference toward using the distal APA sites, which was not present in somatic cell clusters (Fig. 4B). A previous study demonstrated that *ret-1*, which encodes *C. elegans* Reticulon homologue, exhibits two different splicing forms in muscle and intestine^35^. Consistently, we found that *ret-1* in muscle and intestine cell clusters preferentially utilize proximal and distal APA sites, respectively (Fig. 4C, 4D). Moreover, we found that in hypodermis and germline cell clusters, *ret-1* exhibits the opposite preference for two different APA types (Fig. 4C, 4D). The previous study also reported that the splicing preference of *ret-1* in intestine and muscle is reduced with aging^35^. Similarly, we observed that as age increases, the APA site preference of *ret-1* in the intestine shifted from distal to proximal (Fig. 4D), and in the germline, the proportion of cells with the distal APA site also decreased (Fig. 4D). However, the proximal APA site preference in muscle and hypodermis cells was not affected by aging (Fig. 4D).

**Fig. 4.**
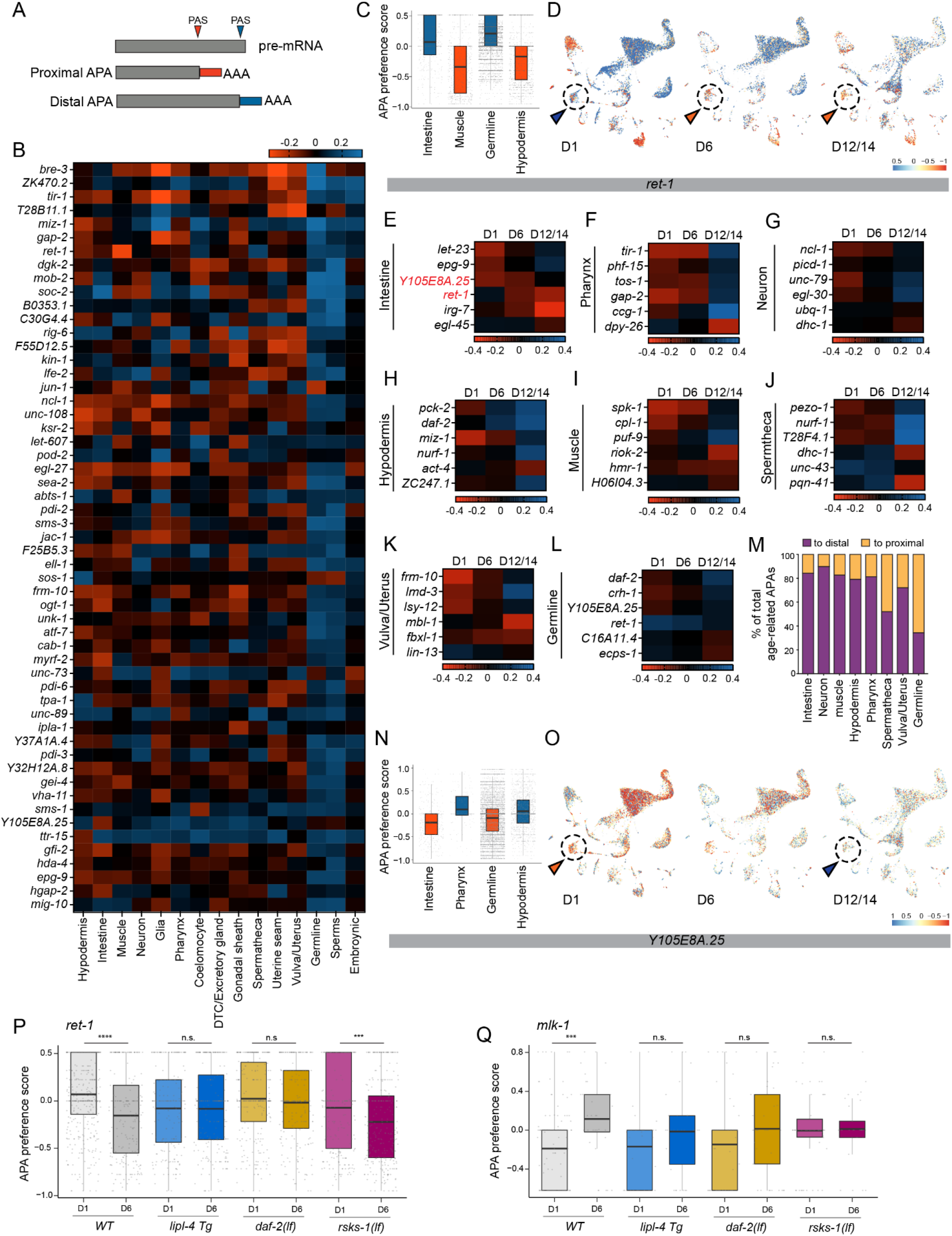
Alternative polyadenylation site usage preference across tissues and ages. (**A**) Schematics of APA site preference towards the proximal or distal polyadenylation site (PAS). (**B**) Heatmaps showing genes with tissue-specific preference for APA sites. (**C**) The APA preference of *ret-1* across four tissues. (**D**) UMAPs showing APA site preference of *ret-1* among all cell types at different ages. Age-related APA changes in the intestine cluster (circle) are highlighted with arrowheads (red for proximal; blue for distal). (**E-L**) For each tissue, six examples of genes exhibiting age-related changes in APA site usage. (**M**) Percentage of genes showing age-related shift towards the distal APA site (purple) or the proximal APA site (orange). (**N**) The APA preference of *Y105E8A.25* across four tissues. (**O**) UMAPs showing APA site preference of *Y105E8A.25* among all cell types at different ages. Age-related APA changes in the intestine cluster (circle) are highlighted with arrowheads (red for proximal; blue for distal). (**P**) The APA site preference shift of *ret-1* in the intestine from day 1 to day 6 suppressed in *lipl-4 Tg* and *daf-2(lf*) but not *rsks-1(lf*) long-lived strains. **** p<0.0001, *** p<0.001, n.s. p>0.05 (**Q**) The APA site preference shift of *mlk-1* in the intestine from day 1 to day 6 was suppressed in all three long-lived strains. **** p<0.0001, n.s. p>0.05

After a systematic search for genes displaying age-related APA changes in different cell clusters, we identified a total of 272 genes in the intestine, 226 genes in neurons, 196 genes in the hypodermis, 133 genes in the muscle, 122 genes in the vulva/uterus, 96 genes in the pharynx, 50 gene in the spermatheca, and 502 genes in the germline (Fig. 4 E-L with 6 examples shown for each tissue, Table S4). This suggests that during aging, various types of somatic and germ cells exhibit changes in APA site preference, and many genes show age-related changes in APA site preference across multiple cell types. Furthermore, we found that in somatic cell clusters including the intestine, muscle, hypodermis, pharynx and neurons, over 80% of the genes exhibiting age-related APA changes decrease their utilization of the proximal APA sites from day 1 to day 12/14 (Fig. 4M). As an example, we identified *Y105E8A.25*, which encodes the *C. elegans* Rho/Rac guanine nucleotide exchange factor, with tissue-specific APA site preference (Fig. 4B, 4N, 4O). During aging, the APA site preference of *Y105E8A.25* changed in intestine, muscle, and germline cell clusters (Fig. 4O, 4E, 4L).

To investigate whether and how pro-longevity mechanisms affect age-related APA changes, we focused on APAs that exhibited significant differences between day 1 and day 6 in WT worms and determined whether their age-related changes were suppressed in long-lived strains. We found that most age-related APA changes in somatic cell types were suppressed by at least one pro-longevity mechanism, with suppression rates of 97.1% in the intestine, 95.4% in neurons, 92.3% in the muscle, 83.4% in the hypodermis, 100% in the pharynx, 96.7% in the vulva/uterus, and 100% in the spermatheca (fig. S5). Furthermore, these age-related APA changes exhibited different suppression patterns in three long-lived strains (fig. S5, Table S5). For example, the shift in the preference from the distal to proximal APA site in intestinal *ret-1* from day 1 to day 6 was suppressed in the *lipl-4 Tg* and the *daf-2(lf*) strains, but not in the *rsks-1(lf*) strain (Fig. 4P); In another example, the age-related shift in the preference from the proximal to distal APA site in intestinal *mlk-1*, which encodes mitogen-activated protein kinase kinase kinase, was fully suppressed in the *rsks-1(lf*) strain, but was only reduced by less than half in the *lipl-4 Tg* and the *daf-2(lf*) strains (Fig. 4Q). These results suggest that APA undergoes age-related changes in different cell types, which can be specifically regulated by different pro-longevity mechanisms.

## Discussion

Together, our studies provide adult aging cell atlases for multicellular organisms, leading to the discovery of transcriptomic signatures and functional features for different cell types in adulthood and during aging. Our machine-learning-based cell-type-specific transcriptomic aging clocks pave the way for understanding the anti-aging effect of different pro-longevity interventions in different cell types and gaining molecular insights into these tissue-specific effects. Lysosomal lipid signaling, IIS and TOR signaling have been examined in this study and more regulatory mechanisms can be investigated in future studies using the open-access platform (http://mengwanglab.org/atlas). Through these analyses, the intestine and neurons emerge as major tissues that are sensitive to age-related changes and exhibit differential responses to different long-lived conditions. As the main site to receive external inputs in *C. elegans*, these two tissues are directly influenced by environmental insults and can produce endocrine signals to dynamically feedback other tissues. Thus, these two tissues stand out as key hubs of longevity regulation.

Reproductive senescence is a hallmark of aging, which disturbs not only reproductive health but also causes somatic dysfunctions through cell-non-autonomous mechanisms. Here we build the first germline-specific trajectory map and transcriptomic aging clock in *C. elegans*. This map allowed us to systematically uncover the molecular characteristics of germ cells at distinctive developmental stages as well as their age-related changes. It is interesting to note that the transcriptomic aging in the germline continues to progress even after reproductive cessation, and the progression is attenuated in the *lipl-4 Tg* and the *daf-2(lf* but not the *rsks-1(lf* long-lived strains. The newly identified genes that are specific to each germ cell stage and undergo age-related changes offer a valuable resource for future studies aimed at understanding the molecular mechanism underlying reproductive aging and its role in somatic aging.

APA is an RNA processing mechanism that plays a crucial role in the control of mRNA metabolism, gene regulation and protein diversification^36^. Our study provides the first systematic profiling of APA changes during aging and its regulation by different pro-longevity mechanisms. Interestingly, APA events exhibit tissue-specific distribution, undergo significant changes during the aging process, and can be differentially regulated by different pro-longevity mechanisms. These tissue-specific events and their regulations may be overlooked in bulk RNA-seq analyses. Interestingly, during aging, somatic cells shift the preference from the proximal to the distal APA site. Previous studies revealed that APA is globally regulated, and the usage of the distal APA site is inversely correlated with the level of core polyadenylation factors^37^. Our finding thus suggests that the level of core polyadenylation factors may decrease with aging in somatic tissues, which can consequently lead to changes in mRNA stability and alternative splicing. Through the open-access platform, future studies could focus on specific APA events to understand their regulatory mechanisms in different cell types and could also systematically explore new APA events under various physiological and pathological conditions. Our studies offer resources for understanding the diversity of longevity-promoting mechanisms at the tissue-specific level, which will facilitate future development of anti-aging and rejuvenation interventions. We openly share these resources using a user-friendly data portal, which will enable other groups to search for cell-type specific transcriptional signatures at different ages and utilize snRNA-seq for analyzing germline differentiation and maintenance, tissue-specific transcriptomic rejuvenation, and APA site preference.

## Acknowledgments

We thank L. Hart, C. Huang and P. Svay for maintenance support; Patrick Allard and Rio Barrere-Cain for their input in establishing the protocol for nuclei extraction. Some strains were obtained from the Caenorhabditis Genetics Center (CGC), which is funded by NIH Office of Research Infrastructure Programs (P40 OD010440). We also thank WormBase and WormAtlas. This project was supported by the Cytometry and Cell Sorting Core at Baylor College of Medicine with funding from the CPRIT Core Facility Support Award (CPRIT-RP180672), the NIH (CA125123 and RR024574) and the assistance of Joel M. Sederstrom.

## Author contributions

Conceptualization: S.M.G., Y.F., and M.C.W. Methodology: S.M.G., Y.Q., A.S.M., H.L., Y.F., and M.C.W. Resources: S.M.G, Q.Z., Y.L., Y.G., and Y.F. Data analysis and visualization: S.M.G., A.S.M, Y.F, M.C.W Project administration: Y.F., M.C.W. Writing – original draft: S.M.G, Y.F, M.C.W. Writing – review & editing: S.M.G., Y.Q., S.Z., A.S.M., Y.L., H.L., Y.F., M.C.W.

## Supplementary Materials

Materials and Methods

Figs. S1 to S5

Tables S1 to S5

## References and Notes

1. Kimura, K.D., Tissenbaum, H.A., Liu, Y., and Ruvkun, G. (1997). daf-2, an Insulin Receptor-Like Gene That Regulates Longevity and Diapause in Caenorhabditis elegans. Science 277, 942–946. 10.1126/science.277.5328.942.

2. Blüher, M., Kahn, B.B., and Kahn, C.R. (2003). Extended Longevity in Mice Lacking the Insulin Receptor in Adipose Tissue. Science 299, 572–574. 10.1126/science.1078223.

3. Apfeld, J., and Kenyon, C. (1998). Cell Nonautonomy of C. elegans daf-2 Function in the Regulation of Diapause and Life Span. Cell 95, 199–210. 10.1016/s0092-8674(00)81751-1.

4. Tatar, M., Kopelman, A., Epstein, D., Tu, M.-P., Yin, C.-M., and Garofalo, R.S. (2001). A Mutant Drosophila Insulin Receptor Homolog That Extends Life-Span and Impairs Neuroendocrine Function. Science 292, 107–110. 10.1126/science.1057987.

5. Papadopoli, D., Boulay, K., Kazak, L., Pollak, M., Mallette, F.A., Topisirovic, I., and Hulea, L. (2019). mTOR as a central regulator of lifespan and aging. F1000research 8, F1000 Faculty Rev-998. 10.12688/f1000research.17196.1.

6. Zhang, Y., Lanjuin, A., Chowdhury, S.R., Mistry, M., Silva-García, C.G., Weir, H.J., Lee, C.-L., Escoubas, C.C., Tabakovic, E., and Mair, W.B. (2019). Neuronal TORC1 modulates longevity via AMPK and cell nonautonomous regulation of mitochondrial dynamics in C. elegans. Elife 8, e49158. 10.7554/elife.49158.

7. Lu, Y.-X., Regan, J.C., Eßer, J., Drews, L.F., Weinseis, T., Stinn, J., Hahn, O., Miller, R.A., Grönke, S., and Partridge, L. (2021). A TORC1-histone axis regulates chromatin organisation and non-canonical induction of autophagy to ameliorate ageing. Elife 10, e62233. 10.7554/elife.62233.

8. Juricic, P., Lu, Y.-X., Leech, T., Drews, L.F., Paulitz, J., Lu, J., Nespital, T., Azami, S., Regan, J.C., Funk, E., et al. (2022). Long-lasting geroprotection from brief rapamycin treatment in early adulthood by persistently increased intestinal autophagy. Nat Aging 2, 824–836. 10.1038/s43587-022-00278-w.

9. Folick, A., Oakley, H.D., Yu, Y., Armstrong, E.H., Kumari, M., Sanor, L., Moore, D.D., Ortlund, E.A., Zechner, R., and Wang, M.C. (2015). Lysosomal signaling molecules regulate longevity in Caenorhabditis elegans. Science 347, 83–86. 10.1126/science.1258857.

10. Savini, M., Folick, A., Lee, Y.-T., Jin, F., Cuevas, A., Tillman, M.C., Duffy, J.D., Zhao, Q., Neve, I.A., Hu, P.-W., et al. (2022). Lysosome lipid signalling from the periphery to neurons regulates longevity. Nat Cell Biol 24, 906–916. 10.1038/s41556-022-00926-8.

11. Elmentaite, R., Conde, C.D., Yang, L., and Teichmann, S.A. (2022). Single-cell atlases: shared and tissue-specific cell types across human organs. Nat Rev Genet 23, 395–410. 10.1038/s41576-022-00449-w.

12. Zeisel, A., Hochgerner, H., Lönnerberg, P., Johnsson, A., Memic, F., Zwan, J. van der, Häring, M., Braun, E., Borm, L.E., Manno, G.L., et al. (2018). Molecular Architecture of the Mouse Nervous System. Cell 174, 999–1014.e22. 10.1016/j.cell.2018.06.021.

13. Regev, A., Teichmann, S.A., Lander, E.S., Amit, I., Benoist, C., Birney, E., Bodenmiller, B., Campbell, P., Carninci, P., Clatworthy, M., et al. (2017). The Human Cell Atlas. Elife 6, e27041. 10.7554/elife.27041.

14. Travaglini, K.J., Nabhan, A.N., Penland, L., Sinha, R., Gillich, A., Sit, R.V., Chang, S., Conley, S.D., Mori, Y., Seita, J., et al. (2020). A molecular cell atlas of the human lung from single-cell RNA sequencing. Nature 587, 619–625. 10.1038/s41586-020-2922-4.

15. Taylor, S.R., Santpere, G., Weinreb, A., Barrett, A., Reilly, M.B., Xu, C., Varol, E., Oikonomou, P., Glenwinkel, L., McWhirter, R., et al. (2021). Molecular topography of an entire nervous system. Cell 184, 4329–4347.e23. 10.1016/j.cell.2021.06.023.

16. Cao, J., Packer, J.S., Ramani, V., Cusanovich, D.A., Huynh, C., Daza, R., Qiu, X., Lee, C., Furlan, S.N., Steemers, F.J., et al. (2017). Comprehensive single-cell transcriptional profiling of a multicellular organism. Science 357, 661–667. 10.1126/science.aam8940.

17. Tang, F., Barbacioru, C., Wang, Y., Nordman, E., Lee, C., Xu, N., Wang, X., Bodeau, J., Tuch, B.B., Siddiqui, A., et al. (2009). mRNA-Seq whole-transcriptome analysis of a single cell. Nat Methods 6, 377–382. 10.1038/nmeth.1315.

18. Kaletsky, R., and Murphy, C.T. (2020). Transcriptional Profiling of C. elegans Adult Cells and Tissues with Age. Methods Mol Biology Clifton N J 2144, 177–186. 10.1007/978-1-0716-0592-9_16.

19. Roux, A.E., Yuan, H., Podshivalova, K., Hendrickson, D., Kerr, R., Kenyon, C., and Kelley, D.R. (2022). The complete cell atlas of an aging multicellular organism. Biorxiv, 2022.06.15.496201. 10.1101/2022.06.15.496201.

20. Li, H., Janssens, J., Waegeneer, M.D., Kolluru, S.S., Davie, K., Gardeux, V., Saelens, W., David, F.P.A., Brbić, M., Spanier, K., et al. (2022). Fly Cell Atlas: A single-nucleus transcriptomic atlas of the adult fruit fly. Science 375, eabk2432. 10.1126/science.abk2432.

21. Martin, B.K., Qiu, C., Nichols, E., Phung, M., Green-Gladden, R., Srivatsan, S., Blecher-Gonen, R., Beliveau, B.J., Trapnell, C., Cao, J., et al. (2023). Optimized single-nucleus transcriptional profiling by combinatorial indexing. Nat Protoc 18, 188–207. 10.1038/s41596-022-00752-0.

22. Hobert, O., Glenwinkel, L., and White, J. (2016). Revisiting Neuronal Cell Type Classification in Caenorhabditis elegans. Curr Biol 26, R1197–R1203. 10.1016/j.cub.2016.10.027.

23. Street, K., Risso, D., Fletcher, R.B., Das, D., Ngai, J., Yosef, N., Purdom, E., and Dudoit, S. (2018). Slingshot: cell lineage and pseudotime inference for single-cell transcriptomics. Bmc Genomics 19, 477. 10.1186/s12864-018-4772-0.

24. Bergen, V., Lange, M., Peidli, S., Wolf, F.A., and Theis, F.J. (2020). Generalizing RNA velocity to transient cell states through dynamical modeling. Nat Biotechnol 38, 1408–1414. 10.1038/s41587-020-0591-3.

25. Leal, S.L., Landau, S.M., Bell, R.K., and Jagust, W.J. (2017). Hippocampal activation is associated with longitudinal amyloid accumulation and cognitive decline. Elife 6, e22978. 10.7554/elife.22978.

26. Koelewijn, L., Lancaster, T.M., Linden, D., Dima, D.C., Routley, B.C., Magazzini, L., Barawi, K., Brindley, L., Adams, R., Tansey, K.E., et al. (2019). Oscillatory hyperactivity and hyperconnectivity in young APOE-ε4 carriers and hypoconnectivity in Alzheimer’s disease. Elife 8, e36011. 10.7554/elife.36011.

27. Palop, J.J., Chin, J., Roberson, E.D., Wang, J., Thwin, M.T., Bien-Ly, N., Yoo, J., Ho, K.O., Yu, G.-Q., Kreitzer, A., et al. (2007). Aberrant Excitatory Neuronal Activity and Compensatory Remodeling of Inhibitory Hippocampal Circuits in Mouse Models of Alzheimer’s Disease. Neuron 55, 697–711. 10.1016/j.neuron.2007.07.025.

28. Zullo, J.M., Drake, D., Aron, L., O’Hern, P., Dhamne, S.C., Davidsohn, N., Mao, C.-A., Klein, W.H., Rotenberg, A., Bennett, D.A., et al. (2019). Regulation of lifespan by neural excitation and REST. Nature 574, 359–364. 10.1038/s41586-019-1647-8.

29. Wang, M.C., O’Rourke, E.J., and Ruvkun, G. (2008). Fat Metabolism Links Germline Stem Cells and Longevity in C. elegans. Science 322, 957–960. 10.1126/science.1162011.

30. Murphy, C.T., McCarroll, S.A., Bargmann, C.I., Fraser, A., Kamath, R.S., Ahringer, J., Li, H., and Kenyon, C. (2003). Genes that act downstream of DAF-16 to influence the lifespan of Caenorhabditis elegans. Nature 424, 277–283. 10.1038/nature01789.

31. Pan, K.Z., Palter, J.E., Rogers, A.N., Olsen, A., Chen, D., Lithgow, G.J., and Kapahi, P. (2007). Inhibition of mRNA translation extends lifespan in Caenorhabditis elegans. Aging Cell 6, 111–119. 10.1111/j.1474-9726.2006.00266.x.

32. Kleemann, G.A., and Murphy, C.T. (2009). The endocrine regulation of aging in Caenorhabditis elegans. Mol Cell Endocrinol 299, 51–57. 10.1016/j.mce.2008.10.048.

33. Langfelder, P., and Horvath, S. (2008). WGCNA: an R package for weighted correlation network analysis. Bmc Bioinformatics 9, 559. 10.1186/1471-2105-9-559.

34. Bhadra, M., Howell, P., Dutta, S., Heintz, C., and Mair, W.B. (2020). Alternative splicing in aging and longevity. Hum Genet 139, 357–369. 10.1007/s00439-019-02094-6.

35. Heintz, C., Doktor, T.K., Lanjuin, A., Escoubas, C.C., Zhang, Y., Weir, H.J., Dutta, S., Silva-García, C.G., Bruun, G.H., Morantte, I., et al. (2017). Splicing factor 1 modulates dietary restriction and TORC1 pathway longevity in C. elegans. Nature 541, 102–106. 10.1038/nature20789.

36. Kelemen, O., Convertini, P., Zhang, Z., Wen, Y., Shen, M., Falaleeva, M., and Stamm, S. (2013). Function of alternative splicing. Gene 514, 1–30. 10.1016/j.gene.2012.07.083.

37. Tian, B., and Manley, J.L. (2017). Alternative polyadenylation of mRNA precursors. Nat Rev Mol Cell Bio 18, 18–30. 10.1038/nrm.2016.116.

38. Hao, Y., Hao, S., Andersen-Nissen, E., Mauck, W.M., Zheng, S., Butler, A., Lee, M.J., Wilk, A.J., Darby, C., Zager, M., et al. (2021). Integrated analysis of multimodal single-cell data. Cell 184, 3573–3587.e29. 10.1016/j.cell.2021.04.048.

39. Aran, D., Looney, A.P., Liu, L., Wu, E., Fong, V., Hsu, A., Chak, S., Naikawadi, R.P., Wolters, P.J., Abate, A.R., et al. (2019). Reference-based analysis of lung single-cell sequencing reveals a transitional profibrotic macrophage. Nat Immunol 20, 163–172. 10.1038/s41590-018-0276-y.

40. Aibar, S., González-Blas, C.B., Moerman, T., Huynh-Thu, V.A., Imrichova, H., Hulselmans, G., Rambow, F., Marine, J.-C., Geurts, P., Aerts, J., et al. (2017). SCENIC: single-cell regulatory network inference and clustering. Nat Methods 14, 1083–1086. 10.1038/nmeth.4463.

41. Kimmel, J.C., Yi, N., Roy, M., Hendrickson, D.G., and Kelley, D.R. (2021). Differentiation reveals latent features of aging and an energy barrier in murine myogenesis. Cell Reports 35, 109046. 10.1016/j.celrep.2021.109046.

42. Berge, K.V. den, Bézieux, H.R. de, Street, K., Saelens, W., Cannoodt, R., Saeys, Y., Dudoit, S., and Clement, L. (2020). Trajectory-based differential expression analysis for single-cell sequencing data. Nat Commun 11, 1201. 10.1038/s41467-020-14766-3.

43. Lange, M., Bergen, V., Klein, M., Setty, M., Reuter, B., Bakhti, M., Lickert, H., Ansari, M., Schniering, J., Schiller, H.B., et al. (2022). CellRank for directed single-cell fate mapping. Nat Methods 19, 159–170. 10.1038/s41592-021-01346-6.

44. Gu, Z., Gu, L., Eils, R., Schlesner, M., and Brors, B. (2014). circlize implements and enhances circular visualization in R. Bioinformatics 30, 2811–2812. 10.1093/bioinformatics/btu393.

45. McGinnis, C.S., Murrow, L.M., and Gartner, Z.J. (2019). DoubletFinder: Doublet Detection in Single-Cell RNA Sequencing Data Using Artificial Nearest Neighbors. Cell Syst 8, 329–337.e4. 10.1016/j.cels.2019.03.003.

46. Friedman, J., Hastie, T., and Tibshirani, R. (2010). Regularization Paths for Generalized Linear Models via Coordinate Descent. J Stat Softw 33. 10.18637/jss.v033.i01.

47. Morabito, S., Reese, F., Rahimzadeh, N., Miyoshi, E., and Swarup, V. (2022). High dimensional co-expression networks enable discovery of transcriptomic drivers in complex biological systems. Biorxiv, 2022.09.22.509094. 10.1101/2022.09.22.509094.

48. Wickham, H. (2011). ggplot2. Wiley Interdiscip Rev Comput Statistics 3, 180–185. 10.1002/wics.147.

49. Chen, E.Y., Tan, C.M., Kou, Y., Duan, Q., Wang, Z., Meirelles, G.V., Clark, N.R., and Ma’ayan, A. (2013). Enrichr: interactive and collaborative HTML5 gene list enrichment analysis tool. Bmc Bioinformatics 14, 128. 10.1186/1471-2105-14-128.

50. Kuleshov, M.V., Jones, M.R., Rouillard, A.D., Fernandez, N.F., Duan, Q., Wang, Z., Koplev, S., Jenkins, S.L., Jagodnik, K.M., Lachmann, A., et al. (2016). Enrichr: a comprehensive gene set enrichment analysis web server 2016 update. Nucleic Acids Res 44, W90–W97. 10.1093/nar/gkw377.

51. Buckley, M.T., Sun, E.D., George, B.M., Liu, L., Schaum, N., Xu, L., Reyes, J.M., Goodell, M.A., Weissman, I.L., Wyss-Coray, T., et al. (2023). Cell-type-specific aging clocks to quantify aging and rejuvenation in neurogenic regions of the brain. Nat Aging 3, 121–137. 10.1038/s43587-022-00335-4.

