## Supplementary figures for "Aging Atlas Reveals Cell-Type-Specific Regulation of Pro-longevity Strategies"

### Supplementary Materials

**This PDF file includes:**

Materials and Methods
Figs. S1 to S5
Tables S1 to S5

#### Materials and Methods

##### *C. elegans*, bacteria strains and maintenance

The following strains were used in this study: N2, CB1370 *daf-2(e1370),* RB1206 *rsks-1(ok1255)*, and MCW14 *raxIs3 [ges-1p::lipl-4::SL2GFP].* The strains N2, CB1370, and RB1206 were obtained from the *Caenorhabditis* Genetics Center (CGC). MCW14 was generated in our lab. All strains were incubated at 20 °C for both maintenance and experiments. *E. coli* OP50 and HT115 were obtained from *Caenorhabditis* Genetics Center (CGC). *E. coli* BW25113, Δlon are from the *E. coli* Keio collection, courtesy of Dr. Christophe Herman (Baylor College of Medicine).

##### Worm preparation

Worms were synchronized using a bleach-based egg isolation method and subsequently starved in the M9 buffer at the L1 stage for 24 hours. Worms of the desired genotype were then seeded to NGM plates pre-seeded with either OP50, BW25113 or Δlon *E. coli*. Approximately 3,000 to 5,000 *C. elegans* were washed off the plates at day 1, day 6, day 12, or day 14. To keep all the cell types including the germline, we used worms that were actively reproducing. For the maintenance of aged worms, we transferred the worm every day by washing them off the plate with M9 buffer and filtering with 40 µm strainers (pluriStrainer, SKU 43-10040-40 and SKU 43-50040-51) to isolate the adult worms while discarding the larvae and eggs in the flow through M9 buffer.

##### Nuclei isolation

Worms were washed three times with PBS and collected in a 1.5 mL tube. We then added 100 µl of homogenization buffer [20] and grind the worms with a pestle motor for 30 seconds on ice. To minimize nuclei adhesion on the surface, all the pestles, tubes and filters were pre-coated with a homogenization buffer or 1x PBS. 900 µl of homogenization buffer was added to wash the pestle, and the total 1mL homogenized sample was transferred into a 1mL Dounce tissue grinder (Wheaton 357538). The grinder was autoclaved overnight in a 220°C oven to deactivate ribonuclease. After placing the grinder on ice, 20 strokes were applied using a loose pestle, followed by another 20 strokes using a tight pestle, while avoiding generating foams. The 1000µl samples were filtered through a cell strainer (35 µm) and then through a Flowmi cell strainer (BelArt, H13680-0040) into a new 1.5 mL tube. The tubes were centrifuged for 10 minutes at 1000 g at 4°C, and the supernatant was removed while not disturbing the pellet, which was hardly visible. The pellet was resuspended with 500 µl 1x PBS with 0.5% BSA and RNAase inhibitor and filtered with a Flowmi cell strainer (BelArt, H13680-0040) again into a 5 mL flow cytometer tube. A total of 20 µl was transferred into another flow cytometer tube and diluted with 180 µl 1x PBS with 0.5% BSA and RNAase inhibitor as an unstained control. Hoechst (Invitrogen 33342) was used to stain the nuclei in 1:1000 working concentration. We coated the 1.5 mL sorting collection tube with 1x PBS with 0.5% BSA and RNAase inhibitor and ~300,000 nuclei were sorted with Hoechst 33342 positive gating, which indicates DNA content, and forward scatter area (FSC-A) gating, representing the particle size above threshold. The intestinal nuclei, which are polypoid (32N), usually form an obvious cluster separating from the other 2N somatic nuclei, and these were also included in our collection. The collection tube was centrifuged at 800g for 8 minutes at 4°C and the sheath buffer supernatant was carefully removed before resuspending the sorted nuclei with 40-50 µl 1x PBS with 0.5% BSA and RNAase inhibitor. The concentration and morphology of the nuclei were checked under a microscope to ensure high-quality nuclei isolation. If the results were desirable, we proceeded to generate gel emulsion with 10X Chromium Controller.

For sorting the nuclei, we used either DB LSR II or Sony MA800 sorter. To obtain the best nuclei morphology and the least RNA degradation, it is recommended to minimize the sorting time. Two researchers could simultaneously set up the sorter and perform nuclei processing.

Single nuclei RNA seq

The nuclei suspensions were loaded onto a 10X Chromium Controller. and library preparation was carried out using the published 10X Chromium Single Cell 3' v2/v3 Solution protocol (Single/dual index). The resulting libraries were sequenced on the HiSeq4000 or NovaSeq6000 platforms, with a depth ranging from 6,306 to 29,862 reads/cell, with the recommended cycle numbers: 26 cycles for Read 1, 8 cycles for i7 index, and 98 cycles for Read 2.

Software utilized for data processing and analysis

| Software/package | Citation |
|---|---|
| Cell Ranger | https://support.10xgenomics.com/single-cell-gene-expression/software |
| Seurat (4.0.5) | [38] |
| SingleR (1.8.1) | [39] |
| AUcell (1.20.1) | [40] |
| scMMD (1.0) | [41] |
| Slingshot (1.8.0) | [23] |
| Tradeseq (1.12.0) | [42] |
| scVelo (0.2.4) | [24] |
| Cellrank (1.5.1) | [43] |
| circlize (0.4.15) | [44] |
| Doubletfinder (2.0) | [45] |
| glmnet (4.1-3) | [46] |
| hdWGCNA (0.2.03) | [47] |
| ggplot2 (3.3.5) | [48] |
| polyApipe (1.0) | https://github.com/MonashBioinformaticsPlatform/polyApipe |
| Prism 9 | https://www.graphpad.com/scientific-software/prism/ |

Single-cell RNA-seq data preprocessing

We used Cell ranger (6.0, 10x Genomics) to align Raw base call (BCL) sequence files or FASTQ files to the *C. elegans* genome (WS282, Wormbase), and generated feature-barcode matrices. Doublets were removed based on the recommended ratio provided by the 10X Genomics Chromium Next GEM Single Cell 3′ Reagent Kits v3.1 user guide (CG000204 Rev D), with a score calculated by Doubletfinder (Github/chris-mcginnis-ucsf). The feature-barcode matrices for each sample were constructed into a Seurat (R package, 4.0.5) object for downstream analysis. Cells were filtered by requiring a minimum of 100 genes to be expressed, and genes were filtered by being expressed in at least 3 cells. Integration of samples was performed with Seurat canonical correlation analysis (CCA) method to remove the batch effects. We tested CCA, reciprocal principal component analysis (PCA) and Harmony (0.1) for integration and chose the method with the best performance. The top 2,000 variable features were used for integration anchor identification. UMAP and tSNE dimension reduction were performed with the first 50 dimensions from PCA. Clustering was performed at multiple Leiden resolutions.

Cell type annotation, marker identification and subclustering

We utilized two approaches to assign cell types to clusters and subclusters. The first approach involved identifying a set of marker genes specific to each cluster using the FindClusters function in Seurat. Then we compared these marker genes to previously reported microscopy-based expression profiles in the literature to confirm the cell type assignment. The second approach utilized the SingleR package to perform automatic annotation by comparing the expression profiles of the cells to those of the reference datasets, enabling accurate annotation of cell types. The reference datasets used in this study include [15,16].

To further dissect the heterogeneity within tissues, we subsetted the Seurat object to include only cells of interest and identified highly variable genes. The data was then subjected to CCA integration and dimensionality reduction followed by clustering to obtain subclusters. The subclusters were then annotated using a similar approach as the main clusters.

Gene set enrichment analysis

The R package AUcell (version 1.20.1) was used to compute the enrichment score of a set of genes in individual single cells. AUCell outputs an enrichment score by using the area under the curve (AUC) to determine whether a critical subset of the input gene set is enriched within the expressed genes for each cell. The AUCell score of a gene set was used to represent the enrichment in single cells. Gene set annotations were extracted from wormEnrichr [49,50], including InterPro classification of protein families (176 gene sets), KEGG pathways networks (111 gene sets), regulatory interactions that connect transcription factors and the genes these factors putatively regulate based on DNA binding site motifs (59 gene sets), and gene ontology libraries for biological process (1711 gene sets).

Cell-type-specific differential expression analysis

We used the Seurat function FindMarkers to perform differential expression analysis between groups of cells. This function employed a non-parametric Wilcoxon rank sum test to identify genes that exhibited differential expression between the groups. Genes with an adjusted *P*-value less than 0.05 were considered to be differentially expressed.

Germline trajectory

To investigate the germline trajectory in our single-cell RNA sequencing data, we performed a series of analyses using various tools. First, germline cells were re-integrated and clustered at a Leiden resolution of 0.5 using Seurat. Two clusters with a sparse UMAP distribution were identified as intra-tissue doublets and removed. Next, we utilized Slingshot with default settings, using the cluster as label inputs and UMAP embeddings as reduced dimension input, to construct cell lineages and recover pseudotime. As a result, we identified two major trajectories and focused our analysis on the trajectory that ended at oocytes.

For the analysis of gene expression patterns along trajectories and for comparing them across genotypes, we applied the figGAM function from the R package TradeSeq. This package uses a generalized additive model (GAM) to fit the expression levels of each gene in each cell and further estimate the non-linear relationship between gene expression and cellular trajectory. By applying this function to our germline trajectory data, we were able to identify key regulatory genes and pathways that control cell fate decisions, leading to a deeper understanding of the biological processes underlying germline development. Overall, our analyses provided valuable insights into the germline trajectory in our sample and the role of different genes and pathways in driving cell fate decisions in the germline lineage.

Velocity/PAGA

To investigate the developmental trajectory of germline cells, we performed a range of analyses using scVelo (0.2.4) and Cellrank (1.5.1). The processed germline Seurat object was converted to an AnnData object, and the top 5000 highly variable genes were selected for velocity calculation. We used the first 30 principal components and set the neighbor number to 50 to perform the velocity analysis. This approach allowed us to gain insights into the changes in gene expression over time and identify the major differentiation pathways and cellular lineages.

In addition to the velocity analysis, we used Cellrank to perform PAGA (Partition-based Graph Abstraction) analysis. This method summarizes the connectivity between the clusters of cells and identifies the major cellular lineages and terminal states. By setting the weight of connectivities to 0.3, we were able to visualize the major cellular lineages and terminal states and gain insights into the underlying biology of the germline cells.

Aging clocks

Aging clocks are machine learning models used to predict age, they were trained on log-normalized expression data from BootstrapCells[51] and true chronological age. R package glmnet (version 4.1-3) was used to fit least absolute shrinkage and selection operator (lasso) regression models via penalized maximum likelihood. Parameters were optimized with 20-fold cross-validation. BootstrapCells were generated based on methods from Buckley *et al.* [51]. Specifically, the transcriptomes of 15 single cells were randomly sampled without replacement from the pool of cells of a given tissue, and gene counts were then summed. This bootstrapping process was repeated 100 times for each tissue. The resulting expression profiles of the BootstrapCells were log-normalized as ln((gene transcripts/cell transcripts)·10,000) and utilized as input for training the aging clocks. Aging clocks are trained on tissues with greater than 50 cells. Aging clock genes were defined as genes with non-zero coefficients in the trained aging clocks, which represent their relative contributions to predicting age in each tissue.

Age-related and tissue-specific GO terms enrichment changes

Tissue-specific gene ontology (GO) terms were determined in WT. For each GO term, the AUCell scores were calculated for all the single cells from WT. The GO terms were assigned to the tissue with the highest median AUCell score. The AUcell scores were then calculated for cells from the same tissue, and median AUCell scores were calculated for day 1, day 6, day 12 and day 14 for the tissue-specific GO terms. Tissue-specific GO terms with a consistent decrease or increase trend during aging were identified.

Tissue-specific age prediction for long-lived strains

We employed tissue-specific aging clocks, which were trained on the expression data of corresponding tissues from WT to predict the biological age of tissues in long-lived strains. To generate the input for the aging clocks, we generated BootstrapCells for each tissue in the long-lived strains. Then log-normalized expression data from BootstrapCells for each tissue were used as input of the trained tissues-specific aging clocks to predict the biological age of tissues for long-lived strains.

scMMD

scMMD is a computational method used to analyze the differences between cell populations in single-cell data. It utilizes the Maximum Mean Discrepancy (MMD) metric to calculate the extent of divergence between two cell populations and a *P*-value. scMMD is an open-source Python package available on GitHub (https://github.com/calico/scmmd).

To perform the analysis, scMMD applies a bootstrap resampling approach with a sample size of 15, running 300 iterations to ensure the robustness and reliability of the results. The method is particularly useful for detecting overall transcriptome differences in gene expression between different cell populations, as it can identify subtle differences that may not be captured by other methods focusing on specific genes.

Co-expression network analysis

We performed co-expression network analysis with weighted gene co-expression network analysis (WGCNA). R package hdWGCNA (version 0.2.03) was used, which is specifically designed to perform co-expression network analysis on single-cell data. We first constructed metacells and selected soft-power thresholds based on the manual guidance. We performed consensus network analysis for each tissue across all the strains. A consensus network can be used to construct a unified network for all the strains and identify networks that are conserved across different genotypes. We subsequently assigned genes to modules based on constructed co-expression networks and set chronological age as a trait to calculate module-trait correlation and identify aging-related modules in different strains. Modules with *P*-value $< 0.001$ and $R^2 > 0.2$ were defined as positively correlated with aging while modules with *P*-value $< 0.001$ and $R^2 < -0.2$ were defined as negatively correlated with aging. Genes in consensus modules that were correlated with aging in at least one strain were extracted. We ran the UMAP algorithm on the hdWGCNA topological overlap matrix (TOM) to visualize these genes in the co-expression networks simultaneously in two dimensions. We subset the columns in the TOM to contain the top one hub genes -- the genes' expressions that are highly connected with other genes -- by eigengene-based connectivity for each module. The organization of each gene in the UMAP space depends on that gene's connectivity with the network's hub genes. We also separately constructed tissue-specific co-expression networks for all the strains to identify genotype-specific aging-related modules. We tested the conservation between all the modules from different genotypes with the Fisher enrich test.

Modules with *P*-values lower than 0.01 are considered conserved modules between genotypes and are linked in circos plot.

Alternative polyadenylation analysis

To analyze alternative polyadenylation (APA) in our 10X Genomics single-cell RNA sequencing dataset, we utilized the R package polyApipe with default settings. We used the *C. elegans* genome (WS282) to generate APA site references for the analysis. The Mann-Whitney-Wilcoxon test was employed to calculate the differences in APA site preference between cell populations, allowing for the identification of statistically significant differences. To identify tissue-specific APA site preference, we used a criterion of FDR-adjusted *P*-value < 0.05 in any inter-tissue comparison (e.g., neuron-muscle) and a gene expression level of at least 20% in the tissue cells. This approach allowed us to identify genes with differential APA site preferences between different tissues at a substantial expression level.

To investigate age-related APA site preference changes in wild-type worms, we combined the day 12 and day 14 samples due to cell number limitations. We then performed a comparison between day 1, day 6, and day 12/14, selecting genes that were expressed in at least 10% of the cells in all ages. This analysis allowed us to identify genes with age-related changes in APA site preference, providing insights into the molecular mechanisms underlying aging in *C. elegans*.

Availability of code and data

We provide an interactive website for exploring all cells, gene expression, subclusters, and APA site preferences. Seurat objects and other analysis outputs can be downloaded from the website resource section.

Lifespan and survival rate measurements

To synchronize the age of worms, a bleach-based egg isolation method was employed, followed by a period of at least 24 hours of starvation in M9 buffer at the L1 developmental stage. To ensure experimental rigor, all genotypes and conditions were tested in parallel. The synchronized L1 worms were then allowed to grow until they reached the first day of adulthood, at which point they were transferred to new plates every two days. In total, around 100 animals were analyzed for each condition and genotype, with 30-50 animals per 6 cm plate. Death was determined by the complete cessation of movement in response to gentle mechanical stimulation. Statistical analyses were conducted using SPSS (IBM software), and the lifespan curve was generated using Prism 9.

#### Supplementary Figures

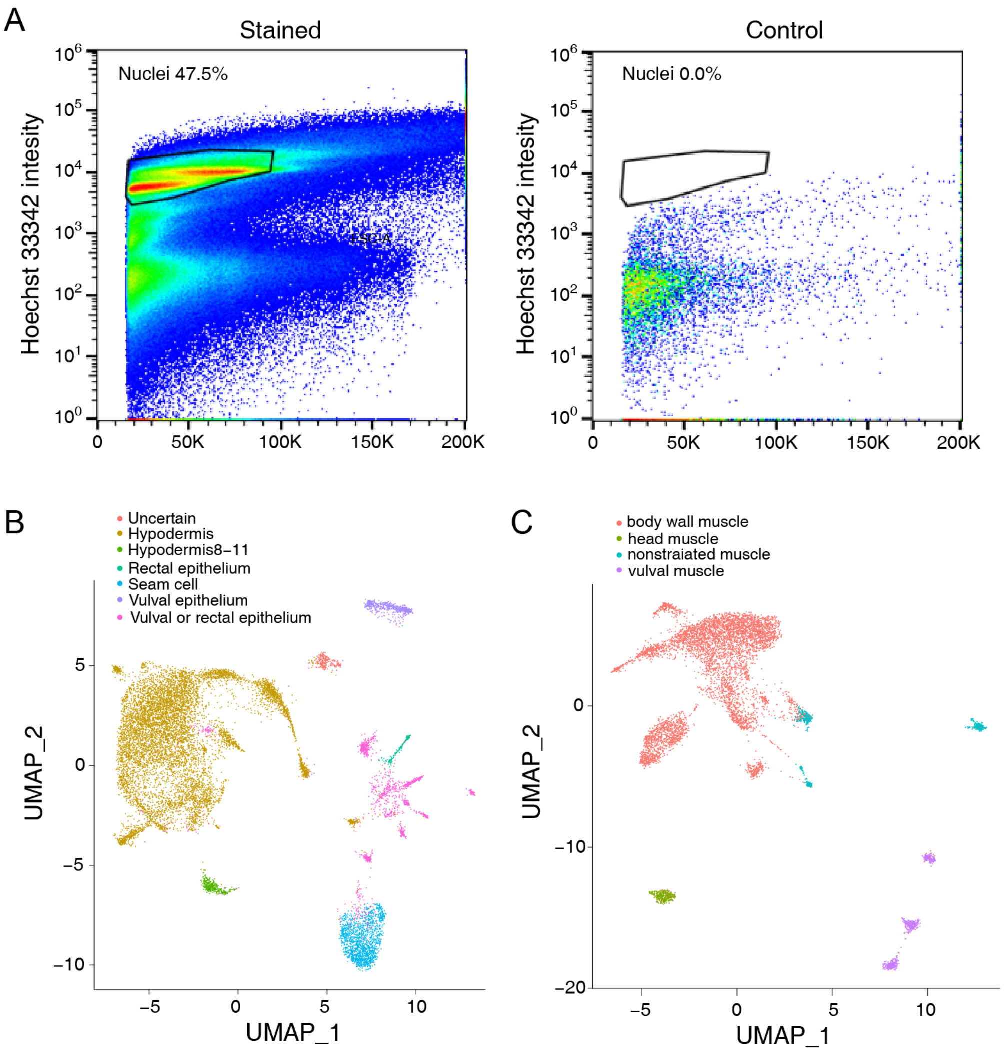


**fig S1. Details of single-nuclei RNAseq and tissue sub-clustering, related to Figure 1**

(A) FACS gating graphs showing a clear separation of nuclei, ensuring the quality of subsequent single-nuclei RNA sequencing analyses. (B) UMAP visualization of hypodermis sub-clustering into seam cell, vulval epithelium, hypodermis cell 8-11, and rectal epithelium. (C) UMAP visualization of muscle sub-clustering into body wall muscle, non-strained muscle, head muscle, and vulva muscle.

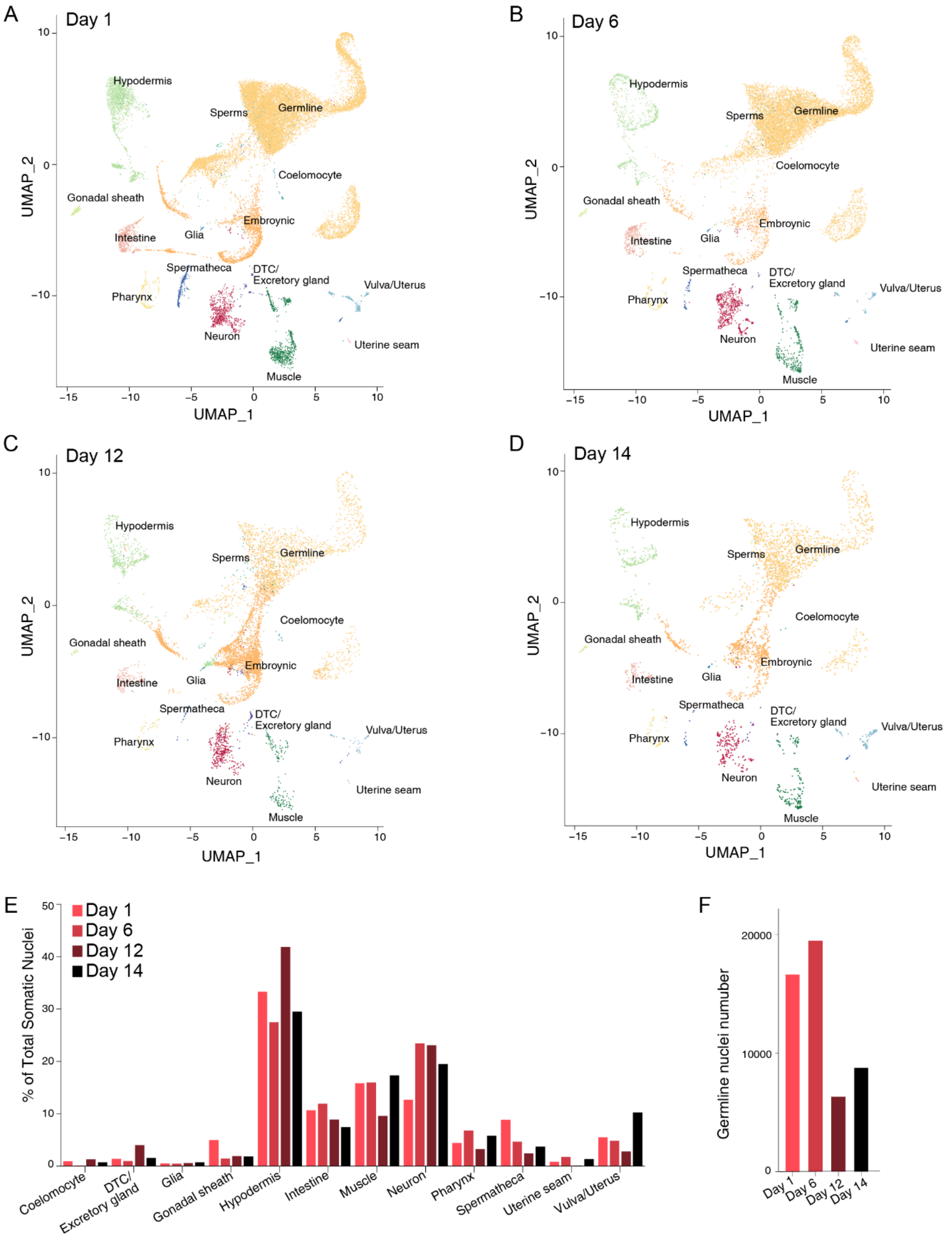


**fig S2. Aging atlases and somatic and germline cell number changes during aging, related to Figure 2**

(A - D) UMAP visualizations of cells from different age groups. (E) Percentage of each cell type in total captured cells at different ages. (F) Numbers of germline nuclei at different ages.

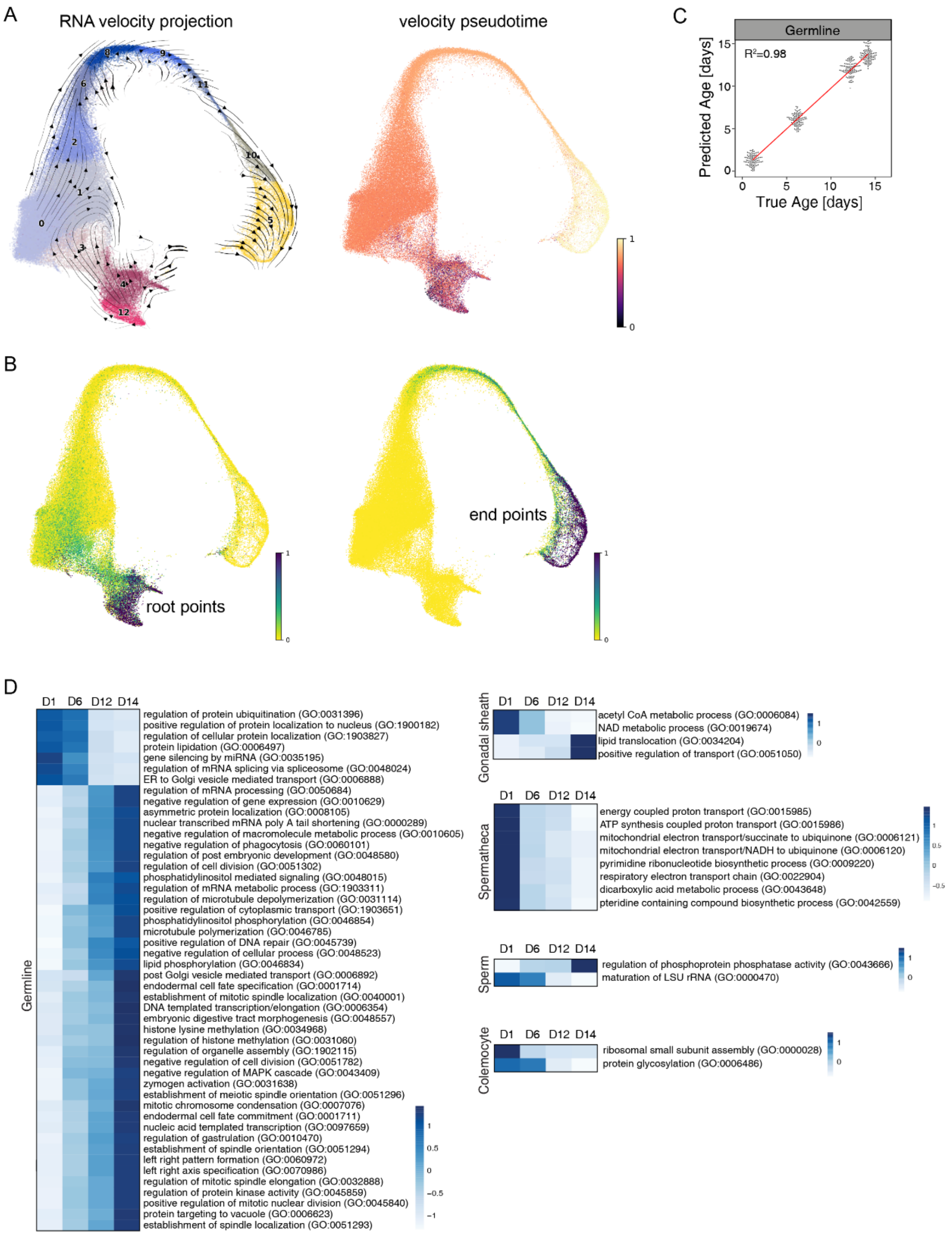


**fig S3. Germline trajectory maps and age-related changes, related to Figure 2**

(A)Velocity projection graph in UMAP embedding and Velocity pseudotime. (B) UMAP graph highlighting predicted root and endpoints that correlated with germline stem cells and oocytes, respectively. (C) Jitter plot showing the correlation between the true age and predicted age from the age clock for the germline. (D) Heatmaps showing changes in the enrichment of GO terms between day 1 and day 6 in WT and long-lived strains for the germline, gonadal sheath cells, spermatheca, sperm and coelomocyte.

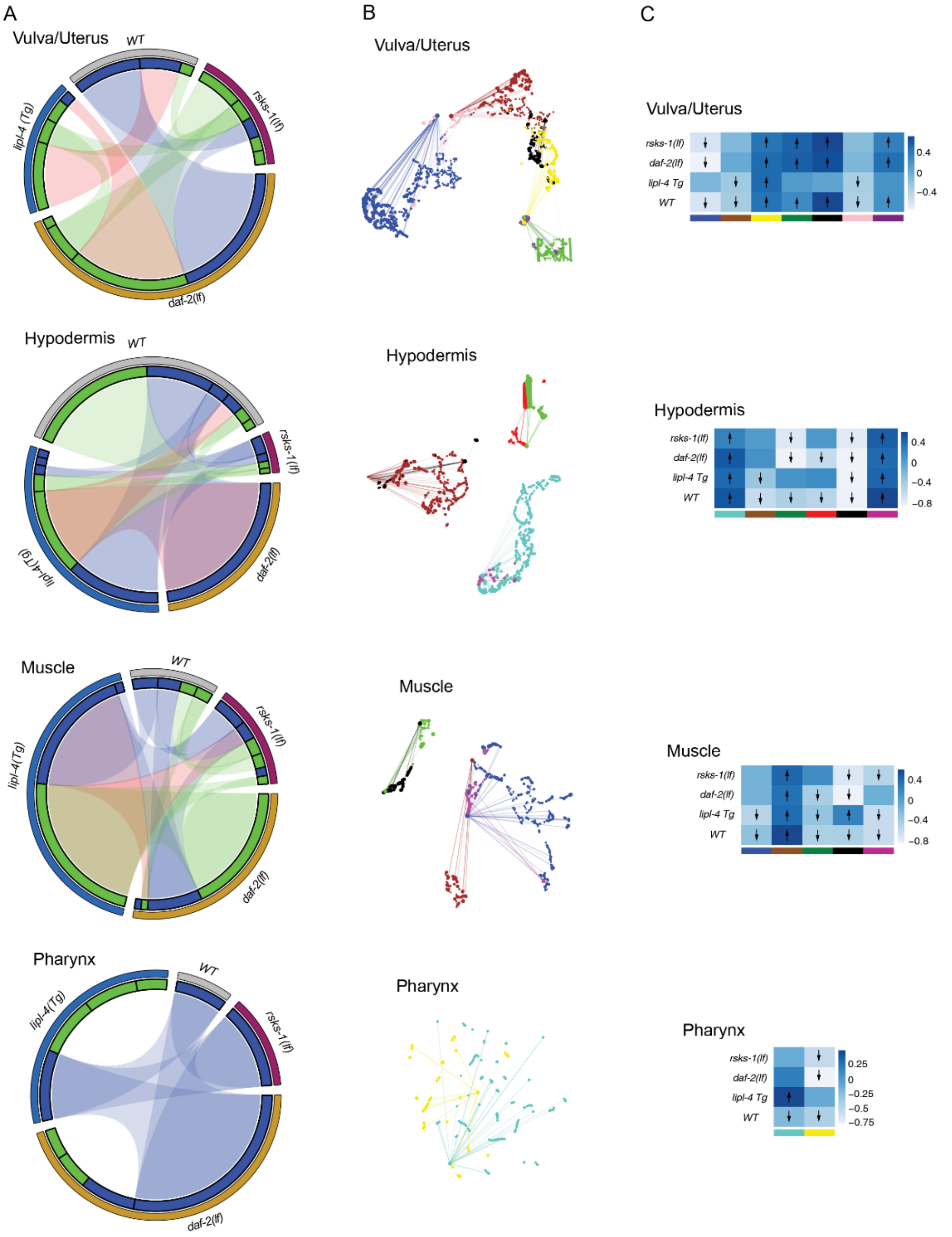


**fig S4. Age-related gene module changes in somatic tissues, related to Figure 3**

(A) Circos plots showing conserved co-expression modules (Fisher's exact test, $P < 0.01$) that were significantly correlated with aging (Pearson's correlation, $P < 0.001$ and $R^2 > 0.2$) in vulva and uterus, hypodermis, muscle, and pharynx between different genotypes. (B) UMAP visualization of the consensus co-expression network for aging-related modules (Pearson's correlation, $P < 0.001$ and $R^2 > 0.2$) in vulva and uterus, hypodermis, muscle, and pharynx. Dots represented genes and were colored by the module they belonged to. Edges represented co-expression between genes. (C) Correlations between consensus co-expression modules with aging, with significant modules (Pearson's correlation, $P < 0.001$ and $R^2 > 0.2$) marked by arrows representing how they correlated with aging.

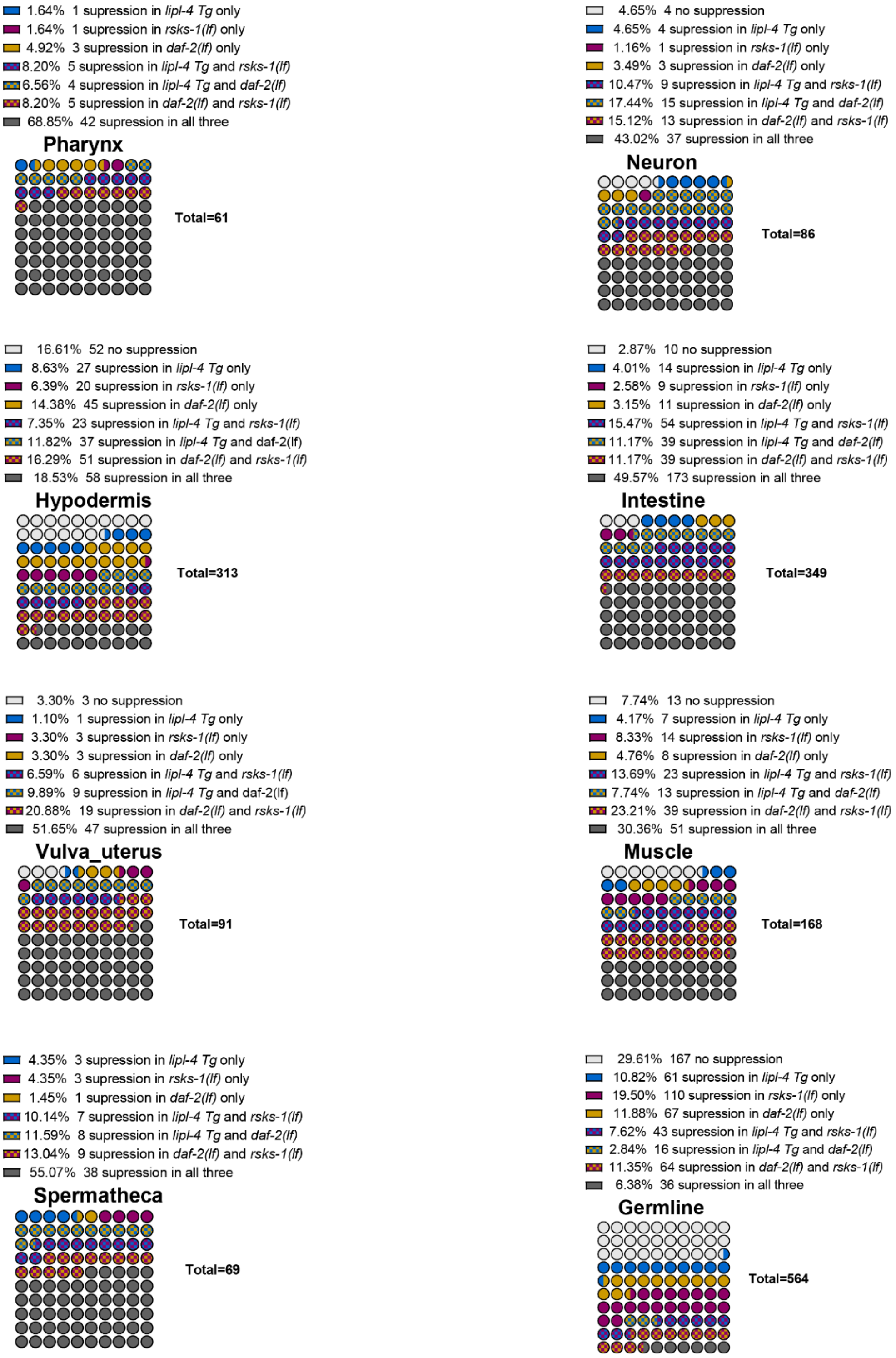


**fig S5. Effects of different pro-longevity mechanisms on age-related APA changes, related to Figure 4**

Waffle plots showing how age-related APA changes are affected by different pro-longevity mechanisms in different tissues.

**Supplementary Table 1. Top 500 genes showing age-related changes in germline temporal patterns.**

**Supplementary Table 2. Differential expressed genes between WT and pro-longevity strains across ages and tissues.**

**Supplementary Table 3. Genes showing tissue-specific preference of APA usage.**

**Supplementary Table 4. Genes showing age-related APA changes in different tissues.**

**Supplementary Table 5. Age-related APA changes suppressed by pro-longevity in various tissues.**
